# Chromatin-Bound PARP1 Correlates with Upregulation of Inflammatory Genes in Response to Long-Term Treatment with Veliparib

**DOI:** 10.1101/2020.03.08.982785

**Authors:** Isabel Alvarado-Cruz, Mariam Mahmoud, Mohammed Khan, Shilin Zhao, Sebastian Oeck, Rithy Meas, Kaylyn Clairmont, Victoria Quintana, Ying Zhu, Angelo Porciuncula, Hailey Wyatt, Shuangge Ma, Yu Shyr, Yong Kong, Patricia M. LoRusso, Daniel Laverty, Zachary D. Nagel, Kurt A. Schalper, Michael Krauthammer, Joann B. Sweasy

**Affiliations:** Department of Cellular and Molecular Medicine, UA College of Medicine, Tucson, AZ; Department of Therapeutic Radiology, Yale University School of Medicine, New Haven, CT; Department of Pathology, Yale University School of Medicine, New Haven, CT; Department of Biostatistics, Vanderbilt University, Nashville, TN; Department of Medical Oncology, West German Cancer Center, University of Duisburg-Essen, Essen, Germany; Department of Biostatistics, Yale University School of Public Health, New Haven, CT; Department of Internal Medicine, Yale University School of Medicine, New Haven, CT; Harvard School of Public Health; Department of Quantitative Biomedicine, University of Zurich, Switzerland

## Abstract

Poly-ADP-ribose polymerase (PARP) inhibitors are active against cells and tumors with defects in homology-directed repair as a result of synthetic lethality. PARP inhibitors have been suggested to act by either catalytic inhibition or by PARP localization in chromatin. In this study, we treat human HCC1937 *BRCA1* mutant and isogenic *BRCA1*-complemented cells for three weeks with veliparib, a PARP inhibitor. We show that long-term treatment with veliparib results in chromatin-bound PARP1 in the *BRCA1* mutant cells, and that this correlates with significant upregulation of inflammatory genes and activation of the cyclic GMP–AMP synthase (cGAS)/ signalling effector stimulator of interferon genes (STING) pathway. In contrast, long-term treatment of isogenic *BRCA1*-complemented cells with veliparib does not result in chromatin-associated PARP or significant upregulation of the inflammatory response. Our results suggest that long-term veliparib treatment may prime *BRCA1* mutant tumors for positive responses to immune checkpoint blockade.

## Introduction

Tumors with high mutation burden (a hypermutator phenotype) are more likely to harbor neoantigens that contribute to immune recognition and response to immunotherapies (for a review see ^1^). Neoantigens originate from somatic mutations that are generated in the tumor (for a comprehensive review see ^2^). Mutant proteins encoded by somatic mutations in tumors are processed by the ubiquitin-proteasome pathway and the resulting mutant peptides are subsequently presented by the major histocompatibility complex class I (MHC class I) to CD8+ T cells or by MHC class II to CD4+ T cells, resulting in the recruitment of these immune cells to the tumor and subsequently leading to tumor death. However, tumors can evade adaptive immune responses through potent tolerogenic mechanisms including the expression of Programmed Cell Death Ligand 1 (PD-L1) in the tumor, which inhibits the proliferation of T cells. Alternatively, tumors presenting Cluster of differentiation (CD) 80 or CD86 may bind the Cytotoxic-T-Lymphocyte-Associated Protein 4 (CTLA-4), acting as an immunosuppressant (Allison 1994; Linsley and Ledbetter 1993; June et al. 1994).

Recent data from a multitude of tumors including melanoma, non-small cell lung cancer (NSCLC), bladder urothelial and microsatellite unstable colorectal carcinomas indicate that the immunogenicity of tumors is prominently influenced by the load of somatic mutations. Higher mutational load, proposed to result in higher levels of neoantigens, correlates with improved responses and progression-free survival in patients treated with the immune checkpoint inhibitors anti-PD-(L)1 or anti-CTLA4 (see for example ^7–9 10,11^).

DNA repair defects induce the accumulation of mutations in the genome, leading to increased levels of neoantigens ^12^. For example, triple negative breast (TNBC) and ovarian epithelial tumors harboring germline *BRCA1, BRCA2* mutations or mutations in genes that function in the homology directed repair pathway (HR) exhibit increased mutational burden and also increased levels of predicted neoantigens ^13,14^. Notably, prominent lymphocyte infiltration is observed in these tumors. Microsatellite unstable colorectal tumors harboring germline mutations in mismatch repair genes also have high mutational burden along with increased levels of predicted neoantigens compared to microsatellite stable tumors ^11^. Importantly, tumors with mismatch repair deficiency displaying high levels of neoantigens respond more favorably to treatment with immune checkpoint inhibitors. A recent case report highlighted the significant response to checkpoint inhibitors in a patient with a metastatic glioblastoma harboring a germline variant of the *POLE* gene ^15^. *POLE* encodes the catalytic subunit of DNA polymerase epsilon, and mutations in the proofreading domain, including the one reported in the cited study, are correlated with an ultra-mutator phenotype and high levels of predicted neoantigens ^15^. The AT-rich interaction domain 1A protein (ARID1A) was recently found to recruit the mutS homolog 2 (MSH2) protein to the chromatin to facilitate mismatch repair ^16^. Importantly, tumors formed from ARID1A-deficient ovarian cancer cells in a syngeneic mouse model displayed increased mutational load along with defective mismatch repair. These tumors regressed in response to treatment with anti-PD-L1, once again demonstrating that high levels of mutations correlate with sensitivity to immune checkpoint blockade^16^. Recently it was discovered that tumors co-mutated in both mismatch and base excision repair genes have increased levels of mutations and predicted neoantigens, and that this correlated with a higher rate of response to treatment with immune checkpoint blockade ^17^. In addition, the clonality of the mutation responsible for neoepitope production is suggested to be an important attribute to elicit responses to immune checkpoint blockade therapy ^18^. Mutational signatures, which reflect the types of mutations produced, rather than their numbers, are also thought to be important for the production of neoantigens, ultimately leading to responses to immunotherapy ^7,19–21^. The DNA repair capabilities of the tumor are likely to be responsible for the majority of the 30 different mutational signatures found in human tumors ^19,20,22^. Therefore, the DNA repair landscape of tumors is an important contributor to the levels of mutations and predicted formation of neoantigens in cancer and is likely to play a significant role in the responses of patients to immunostimulatory therapies. It is important to note that a relatively low mutational load does not preclude tumor response ^7,23^, potentially suggesting that the types of mutations are also important for the generation of specific types of neoantigens that facilitate tumor recognition and elimination.

Moreover, deficiencies in DNA repair may present more opportunities to activate the immune response system ^24^. DNA damage can lead to the production of IFN 1 and other chemokines, which may further activate the immune response and lead to the recruitment of T cells. Breast cancer cells with defects in homologous recombination (HR) have been shown to induce increased levels of cytoplasmic DNA and chemokine release associated with cGAS/STING activation ^25^. Alternatively, severe DNA damage can lead to cell death, and DNA released from apoptotic cells may activate immune response.

In this study, we treated HCC1937 *BRCA1* mutant cells derived from a TNBC and their BRCA1-complemented (BRCA1c) controls with veliparib, a PARP inhibitor, to determine if this treatment induces increased levels of somatic mutations and predicted neoantigens. Veliparib is known to inhibit PARP1 catalytic activity and has been suggested to weakly promote binding of PARP1 to the chromatin ^26–28^. PARP1 recognizes and binds single stranded DNA breaks and catalyzes the production of poly-ADP-ribose chains to facilitate the recruitment of downstream repair enzymes ^29^. Inhibiting PARP1 in *BRCA1* mutant cells may result in the aberrant repair of single- and double-strand breaks, leading to increased mutagenesis ^30–32^. We demonstrate that HCC1937 cells have a high mutational load as compared to BRCA1c cells. Upon treatment with veliparib, BRCA1c cells exhibited an increase in mutations; however, HCC1937 *BRCA1* mutant cells had a similar mutational load as veliparib treated BRCA1c. We did not observe an increase in levels of predicted neoantigens in either cell line. We also show that treatment of HCC1937 *BRCA1* mutant cells with veliparib results in chromatin bound PARP1, and that this correlates with the expression of inflammatory genes and activation of the cGAS/STING pathway.

## Results

### *BRCA1* mutant TNBC cells exhibit higher mutational frequency than BRCA1c cells, but unlike BRCA1c cells, this does not increase with veliparib treatment

HCC1937 cells were previously isolated and cultured from a stage IIB triple negative ductal breast carcinoma and harbor the 5382insC germline mutation in the *BRCA1* gene ^33^. The cells used here also carry the empty pcDNA 3.1 plasmid vector. The presence of the *BRCA1* mutation results in low levels of expression of a truncated BRCA1 protein ^34^, similar to our observations (Figure 1A). In our hands and as previously reported by others ^35–37^, the HCC1937 cells are resistant to PARP inhibitors, in our case veliparib (Figure 1S). This has recently been shown to be due to the absence of the *FAM35A* gene, which comprises part of the REV7-shieldin complex that mediates 53BP1-dependent DNA repair ^38–40^. The HCC1937BRCA1wt (BRCA1c) control cells we employed in our experiments express wild-type (WT) BRCA1 protein expressed from the pcDNA 3.1 plasmid stably integrated into the genome of HCC1937 cells. These cells express full-length BRCA1 protein as previously shown ^34^ and also in our hands (Figure 1A). Importantly, expression of BRCA1 protein in the HCC1937 cells resulted in decreased radiation sensitivity of the HCC1937 BRCA1 mutant cells ^34^. We employed plasmid-based host cell reactivation assays to characterize the ability of these cells to rejoin a double strand break or to conduct homologous recombination^41^. The end joining reporter contains a site-specific double strand break in the 5’-untranslated region of a blue fluorescent protein reporter gene. Fluorescence is abolished unless the double strand break is repaired by end joining, which can occur by the non-homologous end joining (NHEJ) or alternative end joining pathways (alt-NEHJ). We demonstrate that end joining efficiency of a blunt-end double strand break is identical in HCC1937 and BRCA1c cells (Figure 1B lower panel). We also measured homologous recombination efficiency in these cells using both a previously described inter-plasmid recombination assay (Figure 1B upper panel) ^41,42^ and an intra-plasmid recombination assay ^43^ that has been modified to be compatible with other FM-HCR reporter plasmids. We demonstrate that with both assays, homologous recombination efficiency does not differ between HCC1937 and BRCA1c cells. We demonstrate that the BRCA1c cells conduct both homologous recombination and end joining of double strand breaks with equal efficiency as the HCC1937 cells (Figure 1B). In combination, the results suggest that the mutant and WT BRCA1 protein support similar levels of double-strand break repair function in these cells.

**Figure 1.**
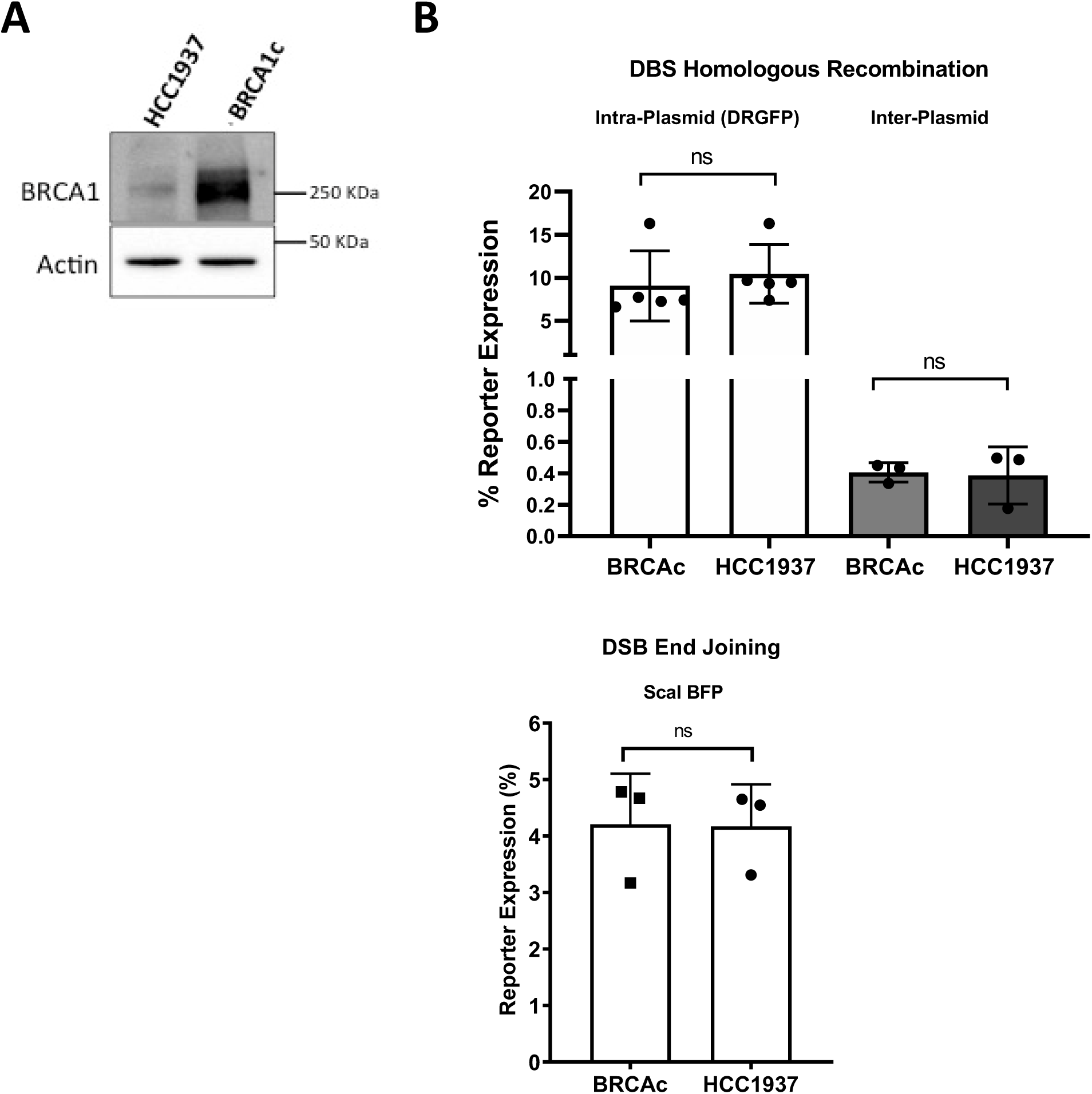
The BRCA1c and HCC1937 cells display similar levels of double-strand break repair. A. Western blot of HCC1937 cells harboring the empty vector (HCC1937) and BRCA1 cells harboring the pcDNA3.1 vector expressing BRCA1 protein (BRCA1c). B. Results of the efficiency of homology directed repair of double strand breaks was similar in *BRCA1* deficient HCC1937 cells and their BRCA1c counterparts. To assess double strand break homologous recombination, cells were transiently transfected with SceI-linearized pDRGFP (open bars), or co-transfected with StuI-linearized pCX-NNX-Δ5GFP and closed circular pCX-NNX-Δ3GFP (shaded bars) upper panel. For double strand break end joining, cells were transfected with Scal BFP (lower panel). For both assays, repair results in expression of GFP.

Next, we determined if treatment of HCC1937 *BRCA1* mutant TNBC cells with veliparib induces increased levels of somatic mutations. Cultures derived from single cell clones of HCC1937 cells and the BRCA1c control cells were treated with 3.6 μM veliparib daily for three weeks. These conditions were chosen to mimic the estimated serum concentration of veliparib used in clinical trials in which patients receive 400 mg BID veliparib for three weeks ^44^. Single cell clones were then isolated, grown to confluence, and interrogated for mutations and predicted neoantigens through analysis of whole genome (WGS), whole exome (WES) and RNA sequencing (RNAseq) (Figure 2A). DMSO was used as the vehicle control for veliparib treatment. Across the exome region, DMSO-treated HCC1937 cells exhibit significantly (p<0.01) elevated numbers of somatic mutations compared to BRCA1c control cells (Figure 2B). Importantly, treatment with veliparib did not increase the numbers of somatic mutations in HCC1937 cells compared to cells treated with DMSO, but there was a statistically insignificant trend toward an increase in the numbers of somatic mutations in BRCA1c control cells treated with veliparib, (p<0.085) (Figure 2B). However, for the cell clones subjected to WGS, we observed a significantly higher number of genome-wide somatic mutations for BRCA1c treated with veliparib compared to DMSO (2408 vs 1118, p < 0.001, test of proportions). For HCC1937, the observed mutations in cells treated with veliparib did not increase compared to DMSO (3774 vs 3958, p=0.98). In accordance with the exome-wide results, the genome-wide data showed significantly higher numbers of somatic mutations for HCC1937 than BRCA1c under DMSO (3958 vs 1118, p < 0.001), corresponding to a somatic mutation frequency of 1.3/MB and 0.4/MB, respectively.

**Figure 2.**
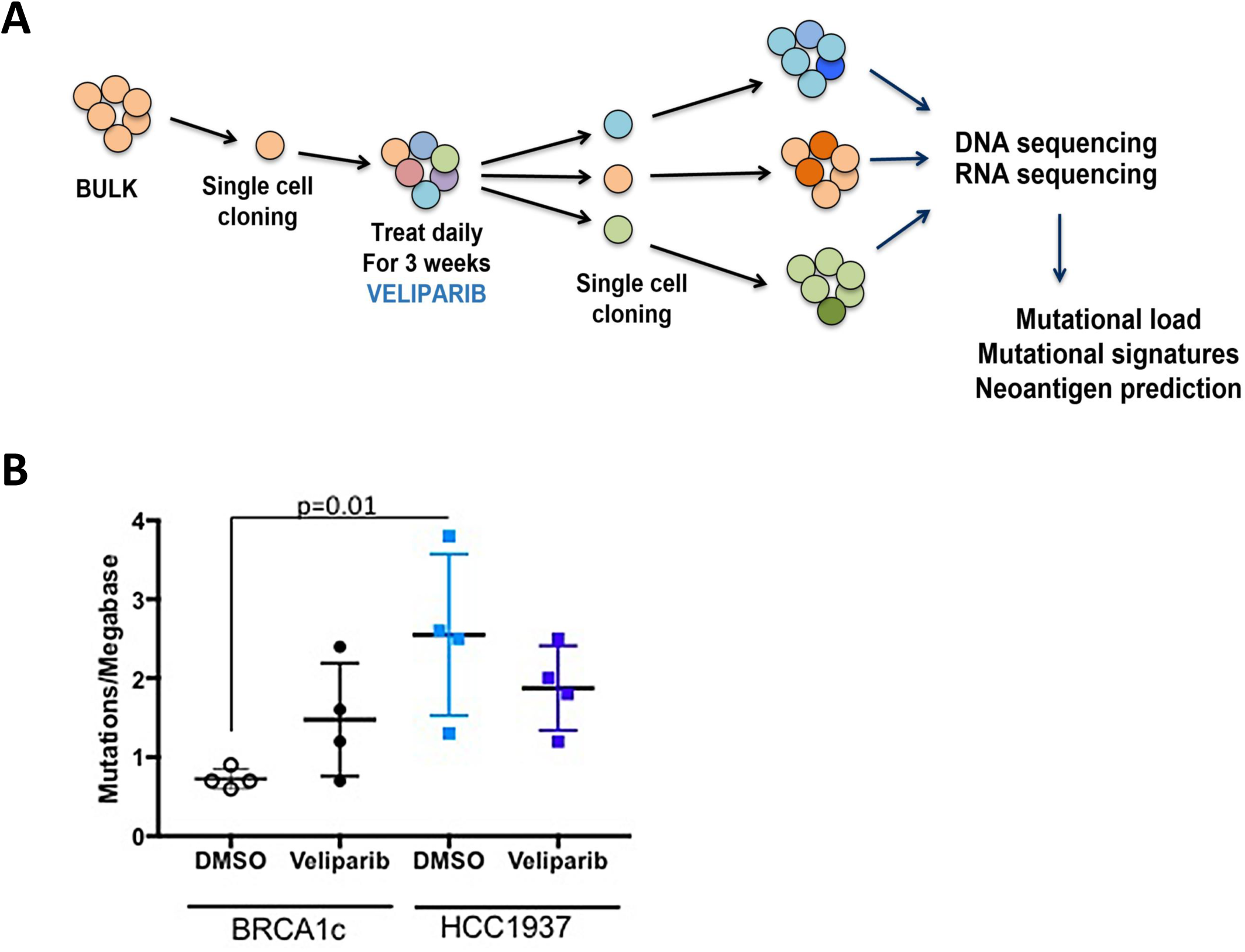
Mutational loads are slightly increased in BRCA1 cells treated with veliparib. A. experimental design. Briefly, single cell clones were treated with veliparib for three weeks. Treated cells were then cloned again, grown to confluence, and sequenced. B. Graph showing mutation frequencies of different clones of BRCA1c cells (black) treated with DMSO (open circles) or veliparib (closed circles) and HCC1937 cells (blue) treated with DMSO (open circles) or veliparib (closed circles).

We also analyzed the mutational signatures of the cell lines after treatment with veliparib ^45^. The overall mutational signature of the HCC1937 cells treated with DMSO is dominated by signature 3, which has been described as a proxy for BRCA1/2 deficiency ^19,20,46^ (Figure 3). The BRCA1c cells treated with DMSO exhibit mutation signature 3, although to a lesser extent than the HCC1937 cells. Therefore, complementation with *BRCA1* seems to have decreased the contribution of mutation signature 3, as would be expected. However, expression of BRCA1 also gives rise predominantly to mutational signature 5, which has been associated with replication stress as a result of the deletion of *FHIT* leading to altered regulation of the *TK1* gene and decreases in nucleotide pools ^47^. Inspection of our RNA sequencing results suggests that both the HCC1937 and BRCA1c cells express normal and equivalent levels of *FHIT* and *TK1*, so deletion of *FHIT* is an unlikely explanation for the appearance of signature 5 in the BRCA1c cells. Upon treatment of HCC1937 and BRCA1c with veliparib, the mutational signatures remain similar to DMSO-treated cells except for the emergence of signature 12. Therefore, signature 12 is not related to *BRCA1* status, but is related to treatment with veliparib and this signature is characterized by T to C mutations on the transcribed strand. Signature 12 is related to errors in transcription-coupled NER that can arise from oxidative DNA damage. In summary, we conclude that *BRCA1* mutated cells have a high mutation load even in the absence of veliparib treatment. Interestingly, while treatment of HCC1937 BRCA1 mutant cells with veliparib does not increase the numbers of observed somatic mutations, there is an increase in mutations in BRCA1c cells treated with this drug.

**Figure 3.**
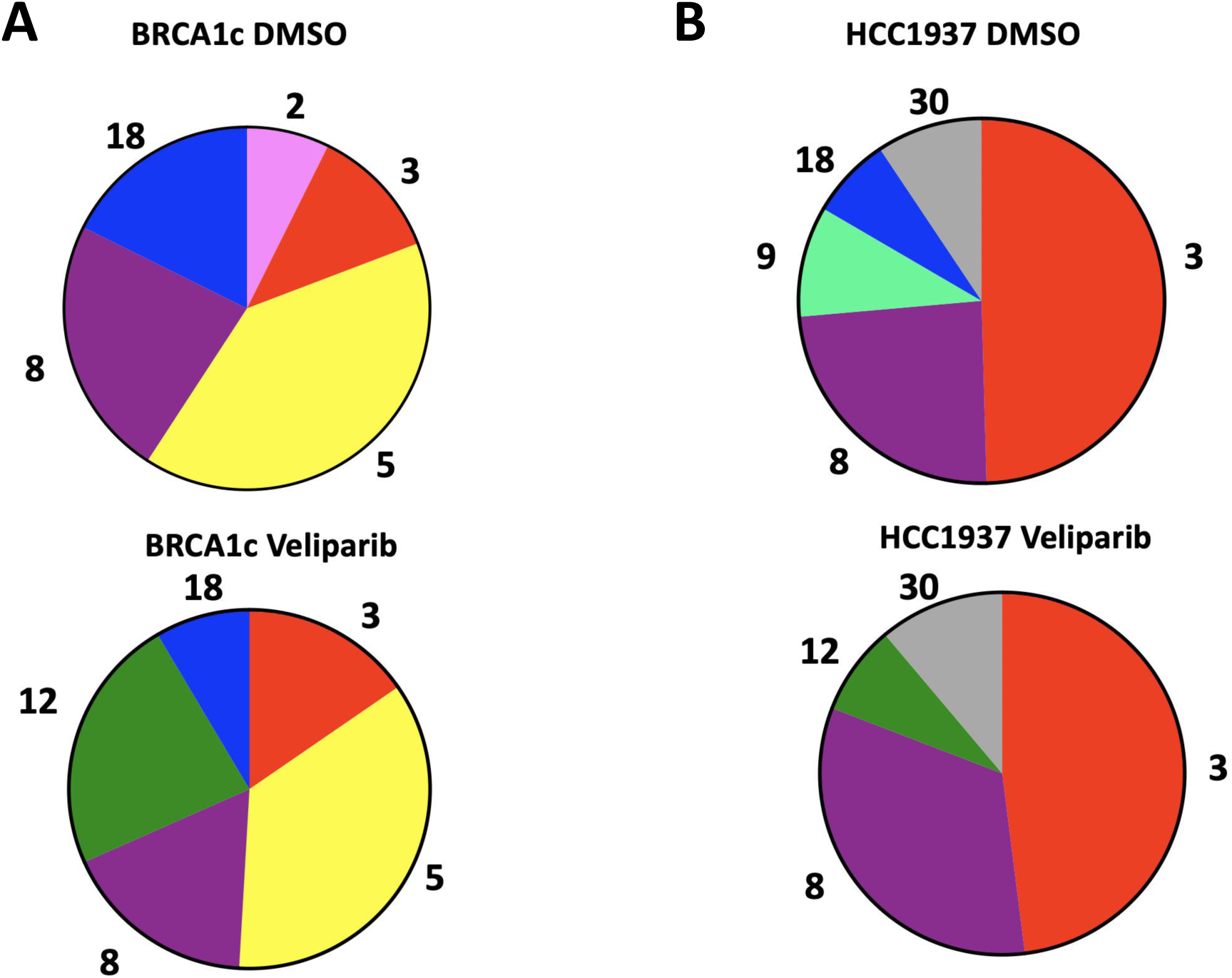
Mutation signatures are not significantly altered upon treatment with veliparib. Shown are pie charts of mutation signature profiles generated as in Methods. Each signature is a different color and labeled with a mutation signature number (see methods); signature 3 (red); 5 (yellow); 8 (purple); 18 (blue); 12 (green); 30 (gray).

### Treatment with veliparib does not significantly increase the numbers of neoantigens

Next, we determined whether treatment with veliparib resulted in potentially increased levels of candidate MHC class I mutant neoantigens, using the pipelines described in Methods. Candidate neoantigens from each condition and identified from these pipelines are listed in Table 1S. Analysis using one-way ANOVA suggests that the numbers of total predicted and expressed neoantigens are greater in HCC1937 cells versus BRCA1c cells treated with DMSO in a manner approaching significance (p=0.057). This is not surprising given that the number of somatic mutations is significantly higher in HCC1937 versus BRCA1c cells. However, treatment of HCC1937 or BRCAc cells with veliparib did not produce significantly increased numbers of predicted neoantigens.

### Veliparib treatment increases expression of inflammatory genes

RNA sequencing data were analyzed to characterize the changes, if any, in the expression of hallmark molecular signature pathways ^48^ in cells treated with veliparib. GSEA (gene set enrichment analysis) reveals that the hallmark pathways that are upregulated in BRCA1c cells treated with veliparib are the interferon α and γ responses (Figure 4A). The interferon α and γ pathways are also upregulated in HCC1937 cells treated with veliparib. In addition, expression of genes in the inflammatory response pathways, TNF alpha and IL6-Jak-STAT3, are also upregulated in HCC1937 cells (Figure 4B). Quantitative RT-PCR was employed to confirm these results. As shown in Figure 4C, CCL5, a member of the TNF alpha pathway; and IRF1, a member of the TNF alpha, Inflammatory Response, Interferon gamma, and IL6-Jak-STAT3 pathways, were significantly upregulated in HCC1937 cells treated with veliparib when compared to BRCA1c cells. CD74, a member of the Inflammatory Response, IFN-alpha and IFN gamma gene sets, and important for antigen presentation, is upregulated in both HCC1937 and BRCA1c cells treated with veliparib, but to a greater extent in BRCAc cells.

**Figure 4.**
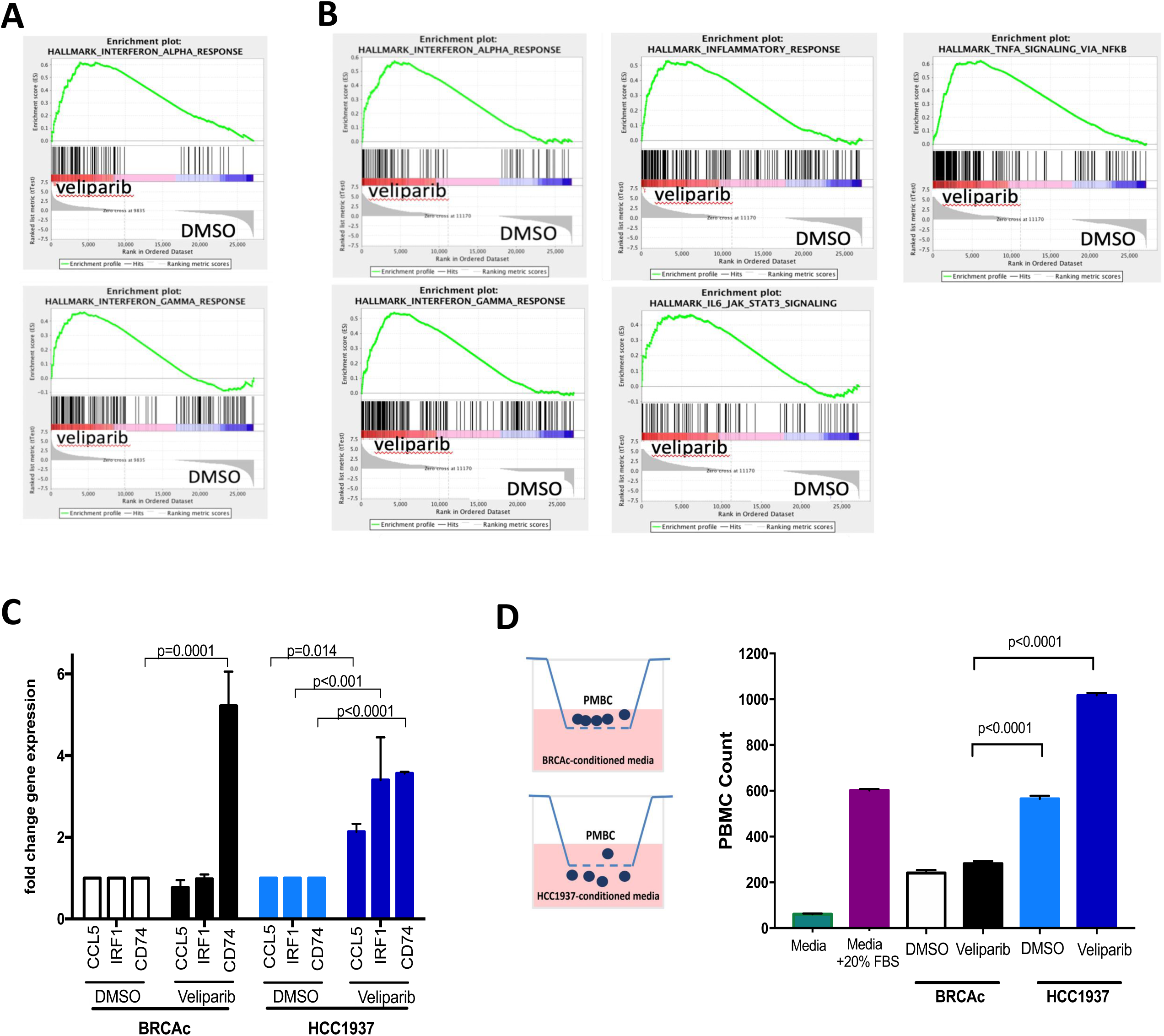
Veliparib treatment induces upregulation of inflammatory pathways and migration of PBMC. A. GSEA hallmark pathways that are upregulated upon treatment of BRCA1c cells with veliparib. Nominal p < 0.001, false discovery rate (FDR) q < 0.001. B. GSEA hallmark pathways that are upregulated upon treatment of HCC1937 cells with veliparib. Nominal p < 0.001, false discovery rate (FDR) q < 0.001. C. Quantitative RT-PCR of specific mRNAs. D. Conditioned medium from culturing either BRCA1c or HCC1937 showed significantly increased numbers of PBMC that migrate towards conditioned media isolated from HCC1937 cells treated with veliparib. The statistical test applied for analysis of significance was the ANOVA with Tukey’s post hoc test.

To confirm that the levels of cytokines increased in cells treated with veliparib, we collected conditioned medium, as described in methods, and asked whether PBMC would migrate towards this medium in response to the cytokines. As shown in Figure 4D, significantly increased numbers of PBMCs migrated towards conditioned medium from HCC1937 versus BRCA1c cells treated with veliparib. This suggests that greater levels of cytokines are present in conditioned medium from HCC1937 veliparib-treated cells in comparison to BRCA1c controls and that they mediate a chemoattractant effect on leukocytes.

### Veliparib activates the STING pathway and results in constitutive activation of the inflammatory response in *BRCA1* deficient cells

Given that the upregulation of CCL5 was observed after treatment of cells with veliparib and that this cytokine is upregulated in response to activation of the cyclic GMP-AMP synthase (cGAS)/stimulator of interferon genes (STING) pathway, we asked if the cGAS/STING pathway was activated. We performed Western blotting analysis to interrogate the phosphorylation status of TBK1 (p-TBK1) and IRF3 (p-IRF3) as markers for activation of the cGAS/STING pathway in cells treated for three weeks with veliparib. Treatment of HCC1937 or BRCA1c cells with veliparib for 3 weeks leads to ∼3-fold increased levels of phosphorylation of TBK1 compared to DMSO-treated controls (Figure 5A). In addition, phosphorylated IRF3 increased in HCC1937 cells after veliparib treatment (Figure 5B). The cGAS/STING pathway has also been shown to crosstalk with the NFκB pathway to induce inflammatory gene expression ^49,50^. Therefore, we interrogated the phosphorylation of NFκBp65 (Ser536) as a marker for NFκB pathway activation. Though both the HCC1937 and BRCAc cells showed an increase in p-NFκBp65, the increase was more pronounced and significant in the HCC1937 cells (Figure 5C). Together these results support the idea that veliparib is inducing a pro-inflammatory state.

**Figure 5.**
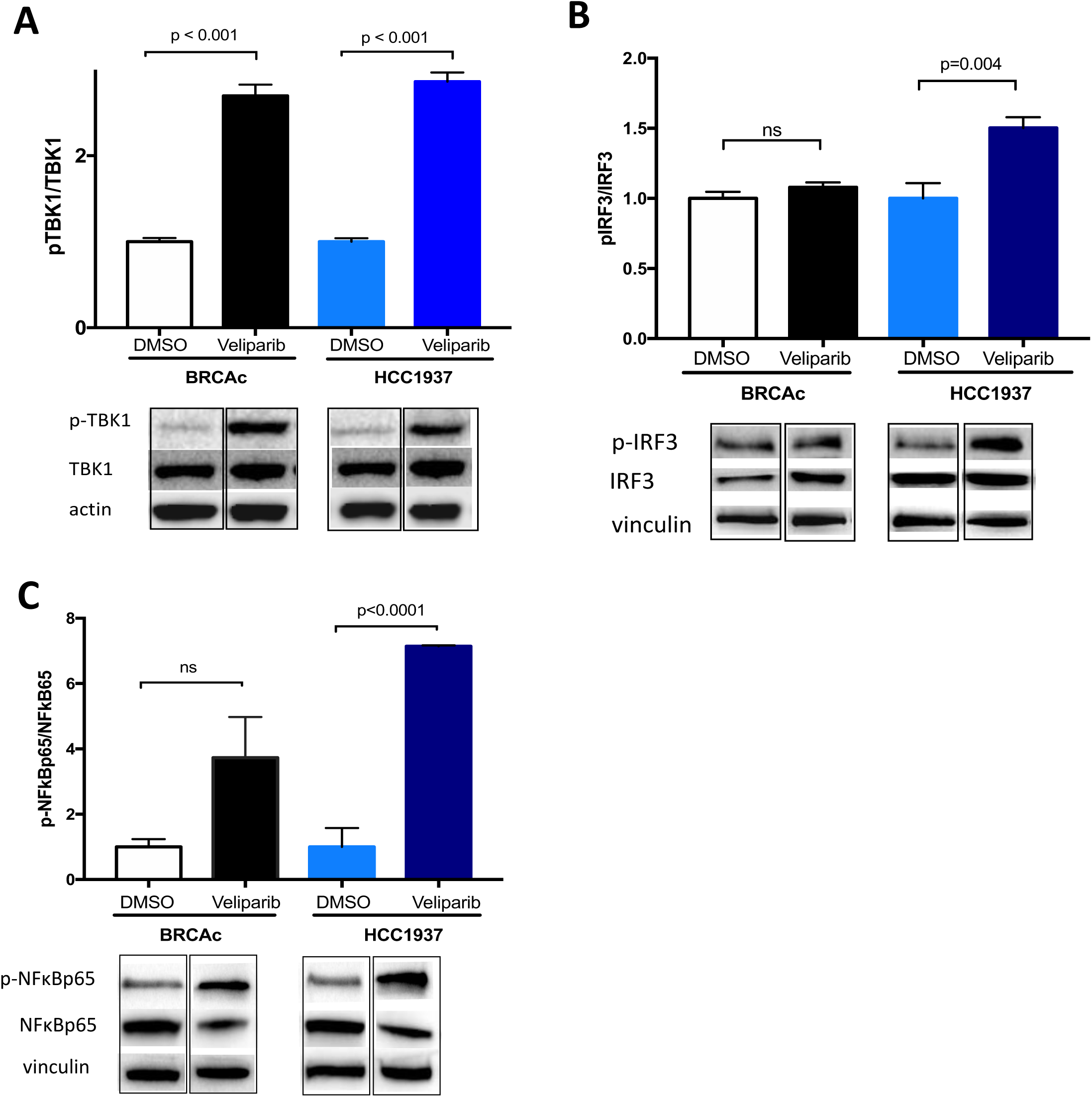
HCC1937 cells have increased phosphorylation of TBK1, IRF3 and NFκBp65 upon treatment with veliparib. Quantification of the ratio of: phosphorylated p-TBK1/TBK1 (A), p-IRF3/IRF3 (B) and p-NFκBp65/NFκBp65 (C) normalized to actin or vinculin. A representative example of a western blot of 3 independent experiments in which cells were harvested after treatment with veliparib for three weeks, as described in methods. Analysis by ANOVA with Tukey’s post hoc test, average of 3 independent experiments.

Activation of the cGAS/STING pathway can result from the presence of DNA in the cytoplasm ^51–53^. Therefore, we examined cells for the presence of cytoplasmic DNA via immunofluorescence. As shown in Figure 6A, there is an increase in cytoplasmic DNA upon veliparib treatment as compared to the DMSO-treated control group for both HCC1937 *BRCA* mutant and BRCA1c cells. However, a difference in the levels of cytoplasmic DNA between HCC1937 *BRCA* mutant and BRCA1c was not observed, which suggests that it is generated as a result of treatment with veliparib and is independent of *BRCA1* status.

**Figure 6.**
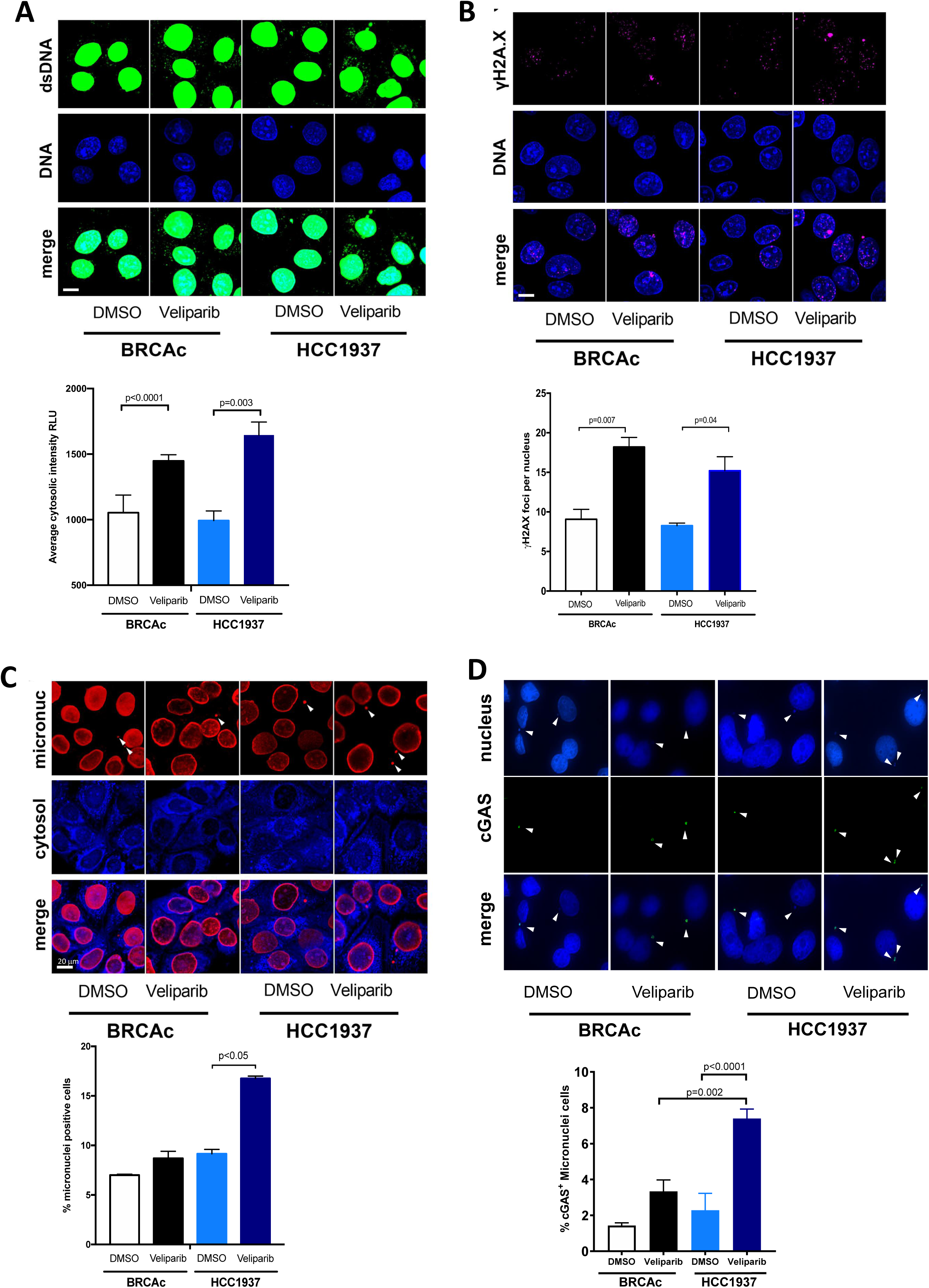
Veliparib promotes cytoplasmic DNA, γH2A.X foci and micronuclei formation after three weeks treatment. A. Quantification and representative images of cytoplasmic DNA immunofluorescence, dsDNA antibody (dsDNA) and DAPI (DNA). B. γH2A.X foci quantification and representative images. C. Micronuclei quantification and representative images. dsDNA antibody was used to stain micronuclei and the cytosol was stained with CellTrace™ cytosol as was described in the methods section. Average of 3 independent experiments was considered and 100 cells per experiments were quantified. Bar= 20 µm

Cytoplasmic DNA can arise as a result of DNA damage, and this has been shown to depend on the activity of the DNA structure-dependent endonuclease Mus81 ^54^. In order to quantify DNA damage, the formation of γH2AX foci was monitored as an indicator of DNA damage. The levels of γH2AX foci increased as a result of treatment with veliparib, although the increase was to a similar extent and independent of the *BRCA* status (Figure 6B).

DNA damage can also result in the formation of micronuclei that provoke the activation of the cGAS/STING pathway ^55,56^. The percentage of cells with micronuclei increased upon veliparib treatment for both cell lines (Figure 6C). However, the percentage of cells with micronuclei in the HCC1937 *BRCA1* deficient cells was significantly higher when compared to the BRCA1c cells. We also show that cGAS is present in micronuclei (Figure 6D). As cGAS recognizes micronuclei, the elevated frequency of micronuclei could lead to the activation of the cGAS/STING pathway in HCC1937 *BRCA1* deficient cells.

### PARP1 becomes chromatin-bound in HCC1937 cells in response to veliparib treatment

Veliparib is a catalytic inhibitor of PARP1 and is also considered to have a weak ability to trap PARP1 to the chromatin ^26,27^. We used an ELISA to determine if catalytic inhibition of PARP1 occurs in cells after three weeks of veliparib treatment. We demonstrate that veliparib inhibits the catalytic activity of PARP1 in both cell lines when compared to DMSO-treated controls, although stronger inhibition occurs in the HCC1937 cells (Figure 7A). To determine if treatment with veliparib results in increased levels of chromatin-bound PARP, we fractionated cells treated with the drug as described ^27^ and performed quantitative Western blotting experiments as described in methods. As shown in Figure 7B, we observe equivalent levels of chromatin-bound PARP1 in DMSO-treated controls in both BRCA1c and HCC1937 cells, suggesting that both cells lines have a similar capacity to associate PARP1 to the chromatin in the absence of drug. However, treatment of BRCA1c with veliparib for three weeks leads to decreased chromatin-bound PARP1 compared to the DMSO controls. Importantly, treatment of HCC1937 cells with veliparib leads to increased levels of chromatin-bound PARP1 when compared to DMSO controls and to the BRCA1c veliparib-treated cells.

**Figure 7.**
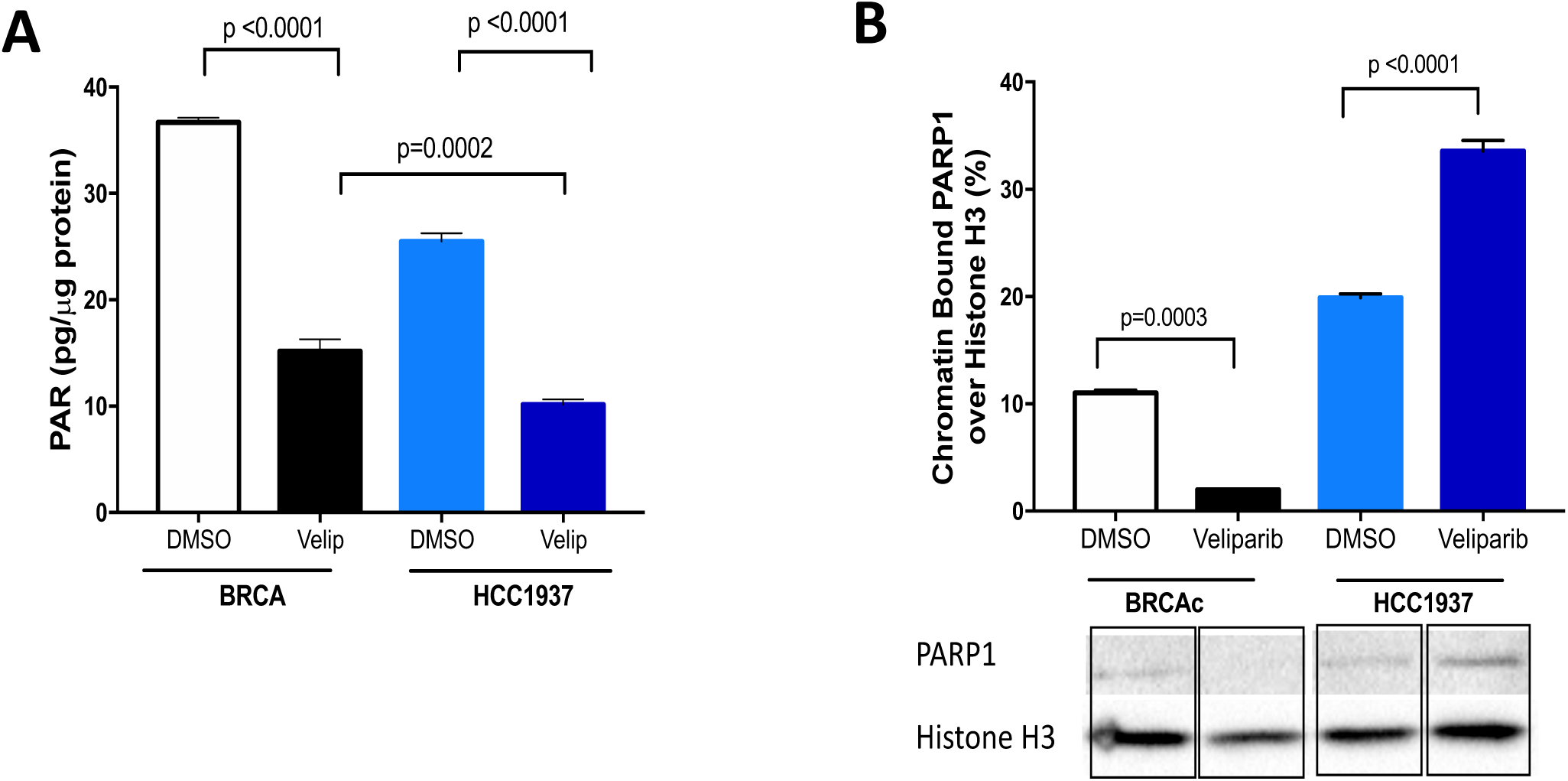
Chromatin-bound PARP1 in HCC1937 and BRCAc cells. A. Quantification of poly-ADP-ribose (PAR) upon treatment of cells with veliparib, using an ELISA assay. B. Quantification of chromatin-bound PARP1 after cellular fractionation and western blotting. ANOVA was used to analyze statistical significance. C. Example of a western blot from a fractionation experiment to quantify chromatin-bound PARP1 in chromatin.

### Chromatin-bound PARP1 is correlated with cytokine expression

To determine if the mechanism of action of veliparib is correlated with mutation frequency, cytokine expression, or migration, we calculated the Spearman coefficients for each of these potential relationships. Interestingly, *CCL5* expression levels are most significantly correlated with chromatin-bound PARP1 (R=0.98; p=0.001) (Table 1). Levels of chromatin-bound PARP1 are also correlated with IRF1 expression levels (R=0.82; p=0.013) and a correlation approaching significance is with migration (R=0.71; p=0.058). Notably, migration is correlated with CD74 expression (R=0.74; p=0.046) and CCL5 expression (R=0.74; p=0.046). In contrast, inhibition of the catalytic activity of PARP1 (PARylation) is negatively correlated with the expression of cytokines and migration. There is not a correlation between PAR or chromatin-bound PARP1 with the numbers of mutations or predicted neoepitopes (not shown).

**Table 1.**

|  | **CCL5** | **IRF1** | **CD74** | **PAR<sup>b</sup>** | **PARP<sup>c</sup>** | **Migration** |
| --- | --- | --- | --- | --- | --- | --- |
| **CCL5** | 1.00 | 0.72 | 0.38 | -0.33 | **0.98** (<0.001) | **0.7** (<0.046) |
| **IRF1** | 0.72 | 1.00 | 0.45 | -0.42 | **0.82** (<0.013) | 0.70 |
| **CD74** | 0.38 | 0.45 | 1.00 | -0.93 | 0.41 | **0.7** (<0.046) |
| **PAR<sup>b</sup>** | -0.33 | -0.47 | 0.40 | 1.00 | -0.38 | -0.76 |
| **PARP<sup>c</sup>** | **0.98** (<0.001) | **0.82** (<0.013) | 0.40 | -0.38 | 1.00 | **0.71** (<0.058) |
| **Migration** | **0.74** (<0.046) | 0.70 | **0.74** (<0.046) | -0.76 | **0.71** (<0.058) | 1.00 |

### Treatment of a TNBC Patient with Veliparib Induces an Inflammatory Response

As a proof of concept and to explore the immunomodulatory effect of veliparib treatment *in vivo*, we studied the immune contexture of baseline and on-treatment tumor samples from a patient with a deleterious germline *BRCA1* mutation (c.4611_4612insG; p.Gln1538Alafs, rs80357915) and advanced TNBC, a biopsy sample was obtained after 6 weeks veliparib treatment 400 mg BID in a phase 2 clinical trial (NCT02849496). As shown in Figure 8A, using multiplexed quantitative immunofluorescence analysis, veliparib treatment induced a 25% increase on CD8+ cytotoxic infiltrating T cells, a 75% increase in CD20+ B-lymphocytes and a 33% reduction in CD4+ helper T cells. Consistent with broader anti-tumor T cell responses, veliparib treatment is also associated with increased TCRα and TCRβ clones (Figure 8B). Individual TCRα and TCRβ clonotypes were also higher, supporting the presence of T cells recognizing additional tumor epitopes after treatment (Figure 8B). Notably, there was a marked increase in TCRγδ clones, suggesting activation of specialized gamma-delta T cell responses. Finally, veliparib is also associated with increased PD-L1 expression in both tumor and immune cells (0% baseline vs 5% tumor-cell IHC score after veliparib and 1 vs 35% immune-cell [IC] IHC score after treatment) (Figure 8C), possibly due to increased adaptive anti-tumor immune pressure and local IFNγ production. Collectively, these results point to a potent immunomodulatory effect of veliparib monotherapy in BRCA1 deficient TNBC. However, additional studies including more cases and clinical outcomes will be required to conclusively assess the consistency and relevance of this finding.

**Figure 8.**
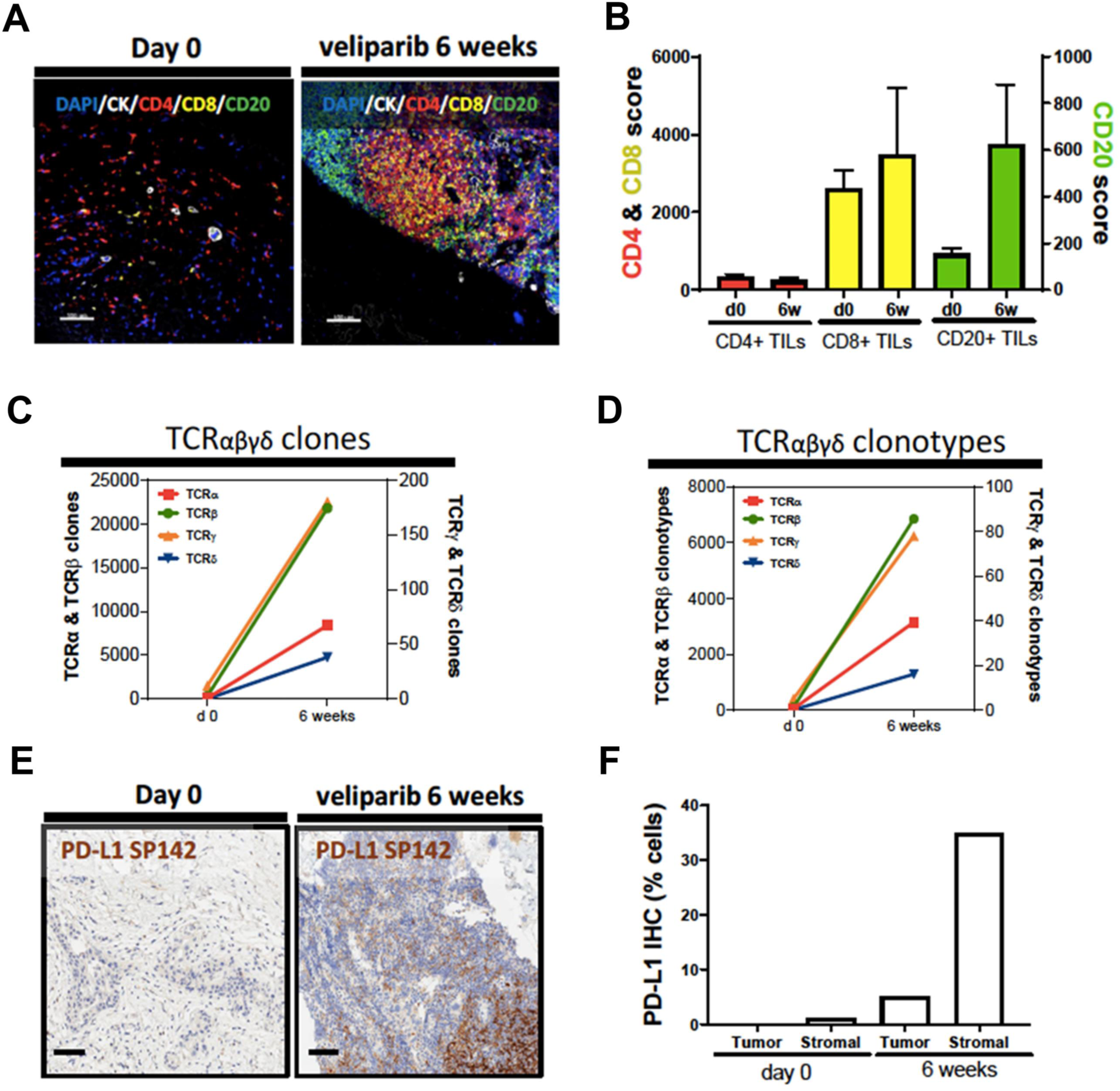
Veliparib induces tumor immune infiltration in a patient with BRCA1 mutant TNBC. A) Multiplexed quantitative immunofluorescence analysis of baseline TNBC tumor samples (day 0) and after 6 weeks of treatment with veliparib (6 weeks). Fixed tumor specimens were simultaneously stained with a panel containing the markers DAPI (blue), cytokeratin (CK, white), CD4 (red), CD8 (yellow) and CD20 (green). Bar=100 μm. The chart indicates the fluorescence level of each immune cell markers before and after treatment. B) Total TCRαβγδ clones (upper panel) and individual clonotypes (lower panel) before and after treatment with veliparib. Each TCR chain is indicated with a different color. C) Chromogenic immunohistochemistry analysis of PD-L1 protein using the antibody clone SP142 in the tumor sample before and after treatment with veliparib. Bar=100 μm. The chart on the right indicate the percentage of membranous PD-L1 positive tumor and stromal (immune) cells as assessed by a pathologist.

## Discussion

In this study we find that HCC1937 *BRCA1* mutant cells exhibit significantly increased numbers of somatic mutations compared to BRCA1c controls, in the absence of veliparib treatment. However, long-term treatment with veliparib induces slightly increased levels of somatic mutations in BRCA1c cells, but not in HCC1937 cells. The lack of induction of mutations in *BRCA1* mutant cells with a PARP inhibitor is consistent with recently published work ^57^. The numbers of predicted neoantigens that are expressed were also higher for HCC1937 cells not treated with veliparib but not in any other case. Interestingly, we find that treatment of our cell lines with veliparib induces upregulation of inflammatory genes, predominantly in the HCC1937 *BRCA1* mutant cells, through the cGAS/STING pathway and that this is related to the levels of cGAS positive micronuclei in the cells. In recent work from other laboratories it has been shown that treatment of *BRCA1*-deficient ovarian tumors with PARP inhibitors in a mouse model induced STING-dependent anti-tumor immunity ^58–60^.

In our system, induction of the inflammatory response correlates with migration of PBMC towards conditioned medium isolated from the HCC1937 *BRCA1* mutant cells supporting a chemoattractant effect on circulating leucocytes. Interestingly, in the one patient we were able to characterize, we observed prominent tumor immune infiltration and expanded T cell clonality after veliparib treatment. Importantly, the upregulation of inflammatory genes along with increased levels of PBMC migration to conditioned medium are correlated with chromatin-bound PARP but not PARP inhibition. Increased mutational load is not correlated with either chromatin-bound PARP, catalytic inhibition or increased proinflammatory signals. This is consistent with recent studies by our group and others showing a lack of correlation between tumor mutational burden and tumor immune infiltration in human solid tumors ^61,62^. These results indicate that tumors with the *BRCA1* 5382insC mutation, and perhaps other *BRCA1* or *2* mutations may respond to immune checkpoint blockers in the absence of treatment with additional agents, given that mutational load is correlated with responses to immunotherapy, and that veliparib may augment the inflammatory response in these cells. Our results also suggest that *BRCA1*-WT cells may respond to treatment with a combination of immune checkpoint blockers and veliparib, since treatment with veliparib induces a slightly increased mutational load and an inflammatory response that differs from that of *BRCA1* mutant cells.

Our results demonstrate that after long-term treatment with veliparib, PARP1 becomes associated within the chromatin of HCC1937 *BRCA1* mutant cells, but very little binding is observed in BRCA1-complemented cells. Recent studies ^63^ indicate that PARP1 binds to and is associated with the DNA via its zinc finger residues, suggesting that chromatin-bound PARP1 can lead to accumulation of stalled replication forks. Although we did not measure the levels of stalled forks directly in this study, the fact that both the HCC1937 and BRCA1c cells are resistant to veliparib, are able to repair double-strand breaks (Figure 1), and exhibit equivalent levels of γH2AX foci (Figure 6B) indicates that stalled forks do not accumulate in either of these cell lines. The absence of the SHLD2 protein in the HCC1937 and BRCA1c cells is likely to result in increased resection and shuttling of DNA repair towards pathways that require resection, including HR and altNHEJ. HCC1937 cells rely on DNA polymerase θ (Pol θ) for survival ^64^, suggesting that altNHEJ may be used to repair the DNA breaks.

The major difference between HCC1937 and BRCA1c cells is that BRCA1c cells express high levels of BRCA1 protein. Our observation that treatment of BRCA1c cells with veliparib leads to decreased levels of chromatin-bound PARP in comparison to cells treated with DMSO or to HCC1937 cells treated with veliparib suggests that the presence of the WT BRCA1 protein in some way prevents or counteracts PARP interaction with chromatin.

Our observation that chromatin-bound PARP1 correlates with upregulation of inflammatory genes, especially in the HCC1937 cells, is consistent with the notion that PARP1 may be associated within the chromatin and exert its effects on transcription. It is documented that PARP1 promotes structural alterations to chromatin, in turn modulating transcription ^67,68^ that is likely taking place through a KDM5B-dependent pathway ^69^. Alternatively, upregulation of inflammation may be a result of chromatin-bound PARP and transient accumulation of blocked replication forks. These forks may rapidly be processed by nucleases, such as MUS81 ^54^, or unrestrained resection, given the absence of *SHLD2* in these cells, leading to the appearance of cytosolic DNA and upregulation of the cGAS/STING pathway. It has previously been shown ^25^ that the upregulation of CCL5 and p-NFκBp65 is a result of STING activation. Consistent with this, we present evidence for upregulation of *CCL5* and induction of p-TBK1, p-IRF-3, and p-NFκBp65, indicators of cGAS/STING pathway activation in HCC1937 cells. Although we also observe induction of p-TBK-1 in the BRCA1c cells treated with veliparib, we do not observe significant upregulation of *CCL5* or induction of phosphorylated-NFκBp65. Importantly, treatment with veliparib induces an increased percentage of cGAS-positive micronuclei in the HCC1937 *BRCA1* mutant cells but <u>not</u> in the BRCA1c cells. It has previously been shown ^55,56^ that recognition of micronuclei by cGAS induces a cellular proinflammatory response. Therefore, we suggest that the presence of micronuclei is important for activation of the robust inflammatory response we observe in the HCC1937 cells. Whether PARP-trapping plays a role in the formation of micronuclei awaits further experimentation.

Expression of the interferon α and γ pathways is induced by treatment with veliparib in the BRCA1c and HCC1937 cells. In addition to this, the hallmark inflammatory, TNFα and IL6-STAT3 pathways are upregulated in HCC1937 cells treated with veliparib. The upregulation of the interferon pathways could lead to activation of cytotoxic lymphocytes in the tumor microenvironment, resulting in tumor cell kill ^70^. This, combined with increased mutational load in the BRCA1c cells, predicts that *BRCA1* non-mutant TNBC may be responsive to combination therapy with veliparib, or a different PARP inhibitor, and an immune checkpoint blocker. Upregulation of genes of the interferon α and γ and hallmark inflammatory pathways in HCC1937 cells indicates that treatment of tumors harboring the *BRCA1* 5382insC mutant with veliparib also induces an inflamed tumor microenvironment. In support of this, we document increased levels of PBMC migration, indicating that the HCC1937 *BRCA1* mutant, but not BRCA1c cells, are secreting factors, most likely chemokines, that attract lymphocytes. In addition, we show enhanced local immune responses upon veliparib treatment in a patient with a deleterious *BRCA1* variant and TNBC. However, veliparib also induces upregulation of anti-inflammatory signals such as TNFα that impairs CD8+ T cells ^71,72^. In fact, blockade of TNFα overcomes resistance to anti-PD-L1 therapy ^71^. In addition, upregulation of the IL6-STAT3 pathway is associated with poor clinical prognosis and suppression of the anti-tumor response (for a recent review see ^73^). Therefore, veliparib treatment of *BRCA1* mutant tumors has the potential to induce both pro-inflammatory and suppressive immune signals that may or may not facilitate responses to immunotherapy treatment. Our results suggest that *BRCA1* status is likely to play an important role in balancing the signals toward the induction of a proinflammatory microenvironment upon treatment with PARP inhibitors.

In summary, we have developed a novel *in vitro* model to assess clinically relevant endpoints of PARPi treatment of TNBC cells. Our results show that *BRCA1* mutant TNBC cells have high mutation burden in the absence of veliparib treatment. In addition, long-term veliparib treatment leads to significant levels of chromatin-bound PARP1 only in the *BRCA1* mutant cells and this correlates with expression of various cytokines, activation of the cGAS/STING pathway, likely through cGAS positive micronuclei, migration of PBMCs and proinflammatory effects in a patient with *BRCA1*-deficient TNBC. Therefore, veliparib treatment of *BRCA1* mutant cells may result in production of T cell inflamed tumors. Treatment of BRCA1c cells with veliparib may lead to increased mutational load, does not result in significant levels of chromatin-bound PARP and migration, but does result in upregulation of the interferon α and β pathways. Increased mutational load and an inflammatory microenvironment are correlated with response to immune checkpoint blockers. Our results suggest that veliparib treatment may prime both *BRCA1* mutant and *BRCA1* wild-type tumors for positive responses to immunotherapy, but perhaps as a result of different underlying mechanisms.

## Methods

### Cell Lines

HCC1937 cells contain the empty vector pcDNA 3.1. BRCA1c cells are HCC1937 cells expressing the BRCA1 protein from pcDNA3.1. Both cell lines were a kind gift from Peter Glazer, Yale University. These cell lines were cultured in IMDM + 15% FBS + 0.15 mg/mL geneticin (G418) antibiotic.

### Samples from a patient with TNBC treated with veliparib

Baseline and on-treatment biopsy samples from a patient with TNBC and a deleterious frameshift mutation in *BRCA1* were prospectively collected. The patient was consented and treated for 6 weeks with veliparib 400 mg bid in the context of the multi-institutional phase 2 clinical trial NCI#10020 (NCT02849496, PI: Patricia LoRusso). The study was open in 2018 and approved by the internal review board (HIC#1608018258/IRB#10020).

### Fluorescence-based Multiplexed Host Cell Reactivation (FM-HCR) assays

Reporter plasmids for double strand break end joining and inter-plasmid homologous recombination were prepared as previously described ^41^. An additional reporter for intra-plasmid homology directed repair was adapted from a previously described DRGFP reporter wherein an engineered ISceI restriction recognition site disrupts GFP expression ^43^. The DRGFP reporter plasmid was obtained from Addgene, expanded in E. coli, and linearized *in vitro* with ISceI. The DRGFP reporter does not express GFP unless homology directed repair of the ISce-induced double strand break restores the wild type sequence using a truncated iGFP fragment donor sequence that lies outside the transcribed region of the plasmid. A control plasmid, “DRGFP-WT” was derived from the DRGFP plasmid by replacing the non-fluorescent ISceI-GFP gene with wild type GFP. To calculate HR, green fluorescent reporter expression from the ISceI-linearized DRGFP was normalized to expression from undamaged DRGFP-WT. HCC1937 and derivative cell lines were maintained below 80% confluence. To measure double strand break repair capacity, cells were trypsinized, suspended in complete medium, and seeded at 125,000 cells per well in a 6-well plate. After 24 hours, a plasmid cocktail containing an end joining reporter (100 ng, blue fluorescent protein reporter gene), a homologous recombination reporter (100 ng, green fluorescent protein reporter gene) and a transfection control plasmid (100 ng, mOrange reporter gene) along with non-fluorescent carrier plasmid (700 ng) was transfected into cells using Lipofectamine 3000 (ThermoFisher) according to the manufacturer’s instructions. Each transfection utilized P3000 reagent (4 μL) and Lipofectamine 3000 reagent (3.75 μL) mixed with the plasmid cocktail in serum-free medium (200 μL, Opti-MEM, ThermoFisher). Cells were incubated for 24 hours after transfection, collected by trypsinization, and analyzed by flow cytometry. Gating and compensation were established by transfection of single color controls and percent reporter expression was calculated as previously described ^42^.

### Treatment of Cells with veliparib

Clonal isolation was performed by serial dilution in 96 well plates. Single clones of HCC1937 and BRCA1c were cultured for 3 weeks in IMDM + 15% FBS + 0.15 mg/mL geneticin (G418) antibiotic in either DMSO or 3.6 µM veliparib (ABT-888), dissolved in DMSO. Cells were seeded at 1:10 in a T-75 flask in IMDM + 15% FBS + 0.15 mg/mL geneticin (G418) antibiotic. The media was aspirated every 24 hrs and the cells were re-treated with fresh media containing either veliparib or DMSO. Once the cells reached 80-90% confluence in the T-75 flask, cells were trypsinized and passaged 1:10 in a new T-75 flask and treated with fresh media 22 containing veliparib or DMSO. After 3 weeks of treatment, clonal isolation was again performed by serial dilution in 96 well plates, followed by expansion into a T-75 flask.

### Isolation of DNA and RNA

5×10^6^ cells were collected and the QIAGEN blood and culture DNA kit (Cat No. 13323) was used per the manufacturer’s protocol. DNA quality was tested with an OD260/280 ratio of 1.8-2. For the purification of total RNA, the QIAGEN RNeasy Plus Mini Kit (Cat No. 74134) was used with a maximum of 1×10^7^ cells and the manufacturer’s protocol was followed. QIAshredder columns (Cat. No. 79654) were used to fully lyse the cells and maximize RNA extraction. RNA quality was determined with OD260/280 ratio of 1.8-2 and an OD260/230 ratio of 1.8-2. RNA integrity was verified by resolving on a 1% agarose gel and we observed a 28s rRNA band twice the intensity of the 18S rRNA band. Extracted DNA and RNA were stored according to manufacturer’s protocol or used immediately for experimentation. RNA was used for RNAseq or for quantitative RT-PCR (qRT-PCR). If used for qRT-PCR the following primers were used:

CXCL10

Forward: GGC CAT CAA GAA TTT ACT GAA AGC A

Reverse: TCT GTG TGG TCC ATC CTT GGA A

CCL5

Forward: CCA GCA GTC GTC TTT GTC AC

Reverse: CTC TGG GTT GGC ACA CAC TT

IRF1

Forward: CTG TGC GAG TGT ACC GGA TG

Reverse: ATC CCC ACA TGA CTT CCT CTT

CD74

Forward: GAT GAC CAG CGC GAC CTT ATC

Reverse: GTG ACT GTC AGT TTG TCC AGC

### DNA and RNA sequencing

Library preparations for whole genome, whole exome, and RNAseq were performed by the Yale Center for Genomic Analysis, Yale University. Whole exome sequencing was performed using the Illumina HiSeq instrument at 140x average coverage (paired end, 100bp). Whole genome sequencing was performed at 22.5x average coverage (paired end,150bp) for the treated clones, and at 66x and 32x for the non-treated HCC1937 cell line and corresponding lymphoblastic cell line, respectively (paired end,150bp). RNA sequencing was performed using the Illunima HiSeq instrument and paired-end sequencing using standard sequencing protocol.

### Analysis of somatic variants and mutational signatures under DMSO and veliparib treatment

We followed best practices in calling variants from our Whole Genome Sequencing (WGS) and Whole Exome Sequencing (WES) data. In short, we used btrim ^74^ to trim ^75^ raw sequencing reads, before subjecting them to bwa alignment, followed by GATK ^76^ processing for calling germline, and MuTect ^77^ processing for calling somatic variants. Variant annotation was achieved using VEP. To derive novel somatic variants under DMSO and veliparib treatment, we used the DMSO treated data as background to identify new variants under veliparib, and the veliparib treated data as background to derive new variants under DMSO. The rationale being that the cells under both conditions are derived from a single clone representing the common variant pool before treatment. Mutational frequencies were derived by tabulating all coding mutations, for both WES and WGS data, in the capture area of the IDT xGen Exome panel and dividing the resulting number by the total capture area (39 MB). Mutational signatures were derived from the WGS data using the deconstructSigs ^45^ R package. For total WGS mutation data, we used a one-sided test of proportions to evaluate whether we see an increased number of mutations in veliparib treated cells, and a two-sided test comparing the number of mutations under DMSO.

### Analysis of RNA-Seq data, GSEA processing

RNA-Seq reads were aligned to the human genome build 38 using STAR (v 2.5.3) ^78^. Subsequently, uniquely mapped reads were summed across all genes in the refseq annotation database using Rsubread feature counts ^79^. Differential expression was analyzed using DESeq in R (v 3.1.2) ^80^. After converting to GenePattern ^81^ GCT format, we preprocessed the raw count data using the GenePattern module PreProcessDataset (version 5.1) with minimum fold change set to 0.5. We then ran the command line GSEA ^82^ tool (version 3.0) with the default setting but for permutation type (set to “phenotype”) and metric for ranking genes (set to “log2_Ratio_of_Classes”).

### Analysis of predicted neoantigens

To evaluate the MHC binding affinity of a given neoepitope, we used NetMHCpan to calculate the predicted binding affinity of a given peptide sequences to specific HLA alleles ^83,84^. Each peptide is a 9-mer sequence as it is the predominant peptide length presented by HLA ^85^. We used the Protein Information Resource (PIR) to determine whether the neoepitope sequence is a non-naturally occurring peptide.

### Chromatin-bound PARP1

After growing for 3 weeks in veliparib, as described above, HCC1937 and BRCA1c cells were grown to confluence. The cells were harvested, washed once with ice cold PBS and centrifuged. From the pellet, subcellular protein fractionation was performed as described in the Subcellular protein fractionation kit for cultured cells (Thermo Scientific; Cat. No. 78840). Fractions were stored in −80°C per manufacturer’s protocol. The next day, PARP1 and histone H3 were detected in the samples by western blot was performed using PVDF membranes as we describe below in WB section. The total intensity of the PARP1 band in the chromatin-bound fraction was divided by the Histone H3 band intensity to obtain % of chromatin-bound PARP1.

### PAR assay using ELISA

This protocol was performed as described ^86^. Briefly, total PAR was quantified using a validated immune-sandwiched assay with denatured cellular extracts.

### Western blotting

The following primary and secondary antibodies were used: TBK1 (108A429 sc-52957; Santa Cruz; 1:1000), p-TBK1 (5483S; Cell Signaling;1,000), IRF3 (11904S; Cell Signaling;1,000), p-IRF3 (37829S; Cell Signaling;1,000), NFκβp65 (8242S; Cell Signaling;1,000), p-NFκβp65 (3033S; Cell Signaling;1,000), β-Actin (A5441; Sigma; 1:5,000), Histone H3 (07-690; Millipore; 1:25000), PARP1 (sc-7150; Santa Cruz,1:500), anti-mouse (NXA931V; GE Healthcare 1:5,000) and anti-rabbit (NA9340; GE Healthcare; 1:25,000). Protein extraction was performed using lysis Buffer (20 mM Tris (pH 8), 200 mM NaCl, 1 mM EDTA, 0.5% NP-40, 10% glycerol). Halt Protease and Phosphatase Inhibitor Cocktail (100x) was added to prevent dephosphorylation. For western blot, the cell lysates plus Laemmli buffer were mixed and heated to 90°C 10 min and loaded in 10% SDS-PAGE gel. Electrophoresis was conducted at 120 V for 1 hr 10 min. Proteins were transferred in 0.2 µm Nitrocellulose Membrane with 20% Methanol for 100 min at 100 V. The membrane was blocked in 5% milk for 1 hr and washed (3×10 min in 1x TBST). Incubation with primary antibody was done overnight to 4°C. After incubation with primary antibody, the blots were washed (3×10 min in 1x TBST) and incubated 1 hr with the adequate secondary antibody. Bands were detected with Thermo ™ SuperSignal™ West Pico and Femto PLUS Chemiluminescent Substrate Catalog: 34080 and 34096.

### Migration assay

Peripheral blood mononuclear cells (PBMCs) from a de-identified patient, were a kind gift from Alfred Bothwell, Yale University, and were obtained using leukophoresis. Using Corning Transwell polycarbonate membrane 5 μm inserts (Sigma Aldrich, St. Louis, MO), 5 × 10^5^ PBMCs were resuspended in Opti-MEM containing 0.5% syringe filtered (0.45 μm syringe filter) BSA and placed in the top chamber of the inserts in a 24-well plate. The PBMCs were incubated for 10 min at 37°C and 5% CO_2_. In the bottom chamber, conditioned media (collected after 3 weeks of treatment of BRCA1c and HCC1937 with either DMSO or Veliparib) was carefully added. On day 3, the transwell insert was removed and a sterile cotton-tipped applicator dipped in sterile PBS was used to remove the remaining cells from the upper chamber carefully without puncturing or damaging the membrane. The transwell insert was then placed into 70% ethanol for 10 min to allow cell fixation. It was then removed, and a cotton-tipped applicator was used to remove the remaining ethanol from the top of the membrane. The transwell membrane was allowed to dry for 10 to 15 min. The membrane was then carefully cut and the numbers of PBMCs that had migrated towards the bottom chamber were quantified using a Cell Titer Glo assay (Promega, Madison, WI) using a standard curve to calculate cell numbers.

### PD-L1 protein detection by immunohistochemistry (IHC)

Formalin fixed and paraffin embedded (FFPE) tissue sections were subjected to PD-L1 staining on a Leica Bond Rx using Leica Refine Polymer Detection Kit as per manufacturer’s instructions. Briefly, 5μm thick sections were baked at 60 °C for 30 min followed by standard dewaxing and rehydration program using Bond Dewax and 100% ethanol. Post rehydration HIER (heat induced antigen retrieval) was performed using ER2 for 20 min at 100 °C and slides were cooled to ambient temperature. Peroxide block was added for 5 min at room temperature followed by primary PD-L1 (0.154 μg/ml, clone SP142, Abcam) for 1 hr at RT. Post Primary reagent was added for 8 min at RT followed by Polymer for 8 min at RT. Mixed DAB Refine was added to slides 2x at ambient temperatures and finally Hematoxylin was added for 5 min at RT. Between each step after peroxide addition, standard Bond Rx washing protocol was applied (3 x Bond wash at ambient temperature). Slides were removed from Leica Bond Rx and subject to manual dehydration and coverslipping. Slides were submerged in ascending alcohol concentrations (70, 80, 90 and 100%[x3] Ethanol) for 3 min each, rinsed in Xylene and submerged in Xylene for 15 min (2x). Slides were coverslipped using Cytoseal mounting media and left to dry overnight. PD-L1 immunostaining was scored by a pathologist (KAS) using light microscopy. Positivity was defined either as the percentage of PD-L1-positive tumor cells (% Tumor or TC score) of any intensity or the proportion of tumor area occupied by PD-L1-positive tumor-infiltrating immune cells of any intensity (% Stromal or IC score).

### Multiplexed Quantitative Immunofluorescence

FFPE tissue sections were stained as previously reported ^87^ using a multiplexed 5-color multiplex immunofluorescence panel containing the markers DAPI for all cells, pancytokeratin for tumor epithelial cells (CK), CD4 for helper T cells, CD8 for cytotoxic T cells and CD20 for B lymphocytes. Briefly, tissue sections were sequentially deparaffinized, rehydrated, and processed for antigen retrieval with EDTA buffer pH 8.0 at 96 °C for 20 min. Slides were incubated with dual endogenous peroxidase block (DAKO) for 10 min at room temperature (RT) and later with 0.3% bovine serum albumin in TBS with 0.05% Tween-20 for 30 min at RT. Primary antibodies were added to the slides in a cocktail for 1 hr at RT to detect the following cells: helper T lymphocytes (anti-CD4 rabbit IgG, 1:100, clone SP35, Spring Biosciences), cytotoxic T lymphocytes (anti-CD8 mouse IgG1κ, 1:250, clone C8/144B, Dako), and B lymphocytes (anti-CD20 mouse IgG2a, 1:150, clone L26, Dako). Tumor epithelial cells were detected using anti-pan-cytokeratin Alexa 488-conjugated mouse IgG1 (1:100, clone EA1/EA3, eBioscience). Secondary antibodies with corresponding fluorophore reagents used were: anti-rabbit EnVision™ (Dako) with biotinylated tyramide (Perkin-Elmer) followed by streptavidin-Alexa 750 conjugate (Perkin-Elmer), anti-mouse IgG1κ (1:100, eBioscience) with Cy3-tyramide (Perkin-Elmer), and anti-mouse IgG2a (1:200, Abcam) with Cy5-tyramide (Perkin-Elmer). Residual horseradish peroxidase activity was quenched in-between secondary antibody/fluorophore incubations using solution of 0.136 g of benzoic hydrazide (0.136 g) in PBS with 50 μl of 30% hydrogen peroxide. Nuclei were counterstained with 2 μg/ml 4’,6-Diamidino-2-Phenylindole (DAPI). Slides were mounted using ProLong Gold antifade reagent (Thermo Scientific). Quantitative measurement of fluorescence in user-defined compartments was obtained using the AQUA method (Navigate BP) as previously described ^87^. Briefly, quantitative immunofluorescence (QIF) scores were obtained for the markers in each fluorescence channel by calculating the marker pixel intensity per compartment (cytokeratin-positive tumor, cytokeratin negative stroma, or in all cells). Scores were normalized by exposure time and bit depth allowing comparison among scores across different samples.

### Multiplexed TCR Sequencing

The multiplexed αβγδTCR analysis using next generation sequencing from FFPE tumor samples was performed as recently reported ^88^. Briefly, total RNA was extracted from FFPE tissue using the AllPrep® DNA/RNA FFPE Kit (Qiagen) with minor modifications. Tumor samples were deparaffinized/rehydrated, then homogenized in 70% ethanol using TissueRuptor II® (Qiagen) at medium setting for 1 min. All other steps were followed according as per manufacturer’s instructions. Total RNA samples were quantified using the Qubit Fluorometer (Thermo Fisher) and normalized by input RNA at 500 ng. Samples were analyzed by TCR sequencing using the Immunoverse assay (Archer Dx), an NGS-ready library preparation that enriches for V(D)J transcripts of the CDR3 region for all four chains of the TCR. Reverse-transcribed cDNA libraries containing the TCR sequences were quantified using KAPA Library Quantification Kit (KAPA Biosystems), and loaded at 4 nM into the MiSeq sequencer (Illumina) v3 chemistry kit spiked with 10% PhiX genome. Data were analyzed using the Archer Analysis online software tool making use of the MiXCR algorithm ^89^. The number of total productive clones and individual clonotypes for each TCR chain were used for the analysis.

### Immunofluorescence staining

The following primary and secondary antibodies were used: anti-γH2A.X (9718 S; Cell Signaling; 1:400), anti-dsDNA (ab27156; Abcam; 1:500), anti-rabbit Alexa Fluor Plus 488 647, anti-rabbit Alexa Fluor Plus 555 and anti-mouse Alexa Fluor Plus 555 (A32731, A32732, A32727; Thermo Fisher Scientific; 1:400) and CellTrace™ (C34554; Thermo Fisher Scientific; 1:500). We seeded 50,000 cells/well in a Millicell^®^ EZ slide (PEZGS0816; Millipore) pretreated with 20 µg/ml collagen. Cells were washed with PBS and incubated with CellTrace™ 20 min at 37°C. After medium removal, cells were fixed with ice-cold methanol at RT 15 min. Each slide was rinse 1x with PBS and permeabilizated with 1% NGS, 0.5% Triton X-100, 8% Sucrose in PBS at RT 20 min. After overnight blocking in 5% normal goat serum with 0.2% Triton X-100 in PBS at 4°C, cells were incubated with primary antibodies in blocking buffer for 1 hr at RT and overnight to 4°C. Following 3 washing steps with PBS + 0.5% Triton X-100, cells were stained with secondary antibodies in blocking buffer for 2 h. DNA was stained with DAPI (1816957; Thermo Scientific Inc; 2.5 μg/mL) for 20 min at RT. Samples were washed twice with PBS + 0.5% Triton X-100 and then rinsed one time with PBS before mounting with DAKO Fluorescence Mounting Medium (S3023; Dako NA Inc.). Images were analyzed with a Nikon Eclipse Ti fluorescence microscope with a Plan Fluor 40 Å∼ /1.30 Oil DIC N2 objective, a CSU-W1 confocal spinning disk unit, an iXon Ultra camera (Andor Technology), MLC 400B laser unit (Agilent Technologies) and NIS Elements 4.30 software (Nikon Corporation). Focinator ^90^, a tool for automated high-throughput analysis was used to analyze the images.

### Statistical analysis

t student and ANOVA analysis with Tukey’s post hoc test were used for statistical analysis with GraphPad Prism 7.0 Software. Errors bars represent standard errors of mean (SEM).

## Author contribution

J.B.S conceived the study and obtained the funding. I.A.C., M.M, M.K., S.O., R.M., K.C., V.Q., D.L., Z.D.N., performed experiments. S.Z., M.K., contributed to data analysis. P.M.L., provided tumor sample. K.A.S., performed tumor multiplexed immunofluorescence and sequencing analysis. J.B.S and I.A.C wrote the paper.

## Acknowledgments

This work was funded by NCI R21 CA216595-01 to JBS and PML;1UM1CA186689 to PML; 2P01CA092584-16 to Z.N.; Rising Tide Foundation For Clinical Cancer Research Grant to PML, a P30 NIH/NCI Immunotherapy Biomarker supplement to PML and KAS; Department of Defense Grant W81XWH-16-1-0160 to KAS and Stand up to Cancer Translational Grants SU2C-AACR-DT17-15 and SU2C-AACR-DT22-17. MM was supported in part by a generous gift from Stephen Sherwin, M.D.

