## Supplementary material for "Chromatin-Bound PARP1 Correlates with Upregulation of Inflammatory Genes in Response to Long-Term Treatment with Veliparib": Supp files

**Supplementary Data**

Figure 1S. HCC1937 and BRCA1c cells are not sensitive to veliparib. Cells were treated with various concentrations of veliparib for 7 days and survival was determined by Cell Titre Glo (Promega) BRCA1c (black); HCC1937 (blue); MCF7 (green).

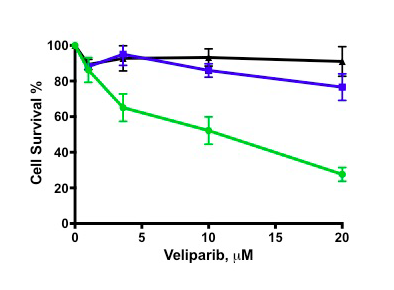
